# High-quality haplotype-resolved genome assembly and annotation of *Malus baccata* ‘Jackii’

**DOI:** 10.1101/2025.07.27.667097

**Authors:** Matthias Pfeifer, Ofere Francis Emeriewen, Henryk Flachowsky, Monika Höfer, Jens Keilwagen, Fang-Shiang Lim, Andreas Peil, Holger Zetzsche, Thomas Wöhner

**Affiliations:** Julius Kühn-Institut (JKI) - Federal Research Centre for Cultivated Plants, Institute for Breeding Research on Fruit Crops, Dresden-Pillnitz, Germany; Institute of Plant Genetics, Department of Molecular Plant Breeding, Leibniz University Hannover, Hannover, Germany; Julius Kühn-Institut (JKI) - Federal Research Centre for Cultivated Plants, Institute for Biosafety in Plant Biotechnology, Quedlinburg, Germany; Julius Kühn-Institut (JKI) - Federal Research Centre for Cultivated Plants, Institute for Resistance Research and Stress Tolerance, Quedlinburg, Germany

## Abstract

*Malus baccata* ‘Jackii’ has been observed to exhibit multiple disease resistances, thus rendering it a promising source for breeding new disease-resistant apple cultivars. Here, we present the first haplotype-resolved genome assembly and annotation of this genotype, achieved by integrating PacBio HiFi sequencing, Hi-C, and mRNA sequencing data with a range of bioinformatic tools and databases. The genome assembly comprises 17 pseudochromosomes with total scaffold lengths of 654.6 Mb and 637.5 Mb for the two haplotypes, respectively. Both haplotypes have scaffold N50 values exceeding 30 Mb, with 42,441 and 46,507 predicted genes, of which 99.9% were successfully annotated. The high quality of this genome is supported by BUSCO analysis values exceeding 97.5% for both haplotypes. This comprehensive dataset is well suited for a wide range of future genomic analyses and is anticipated to benefit apple breeding, particularly in the context of enhancing disease resistance.

## Background & Summary

The apple (*Malus domestica* Borkh.) is among the most popular and important fruits in the world. Different closely related species of apple (*Malus* spp.) can hybridize easily, and the domesticated apple of today contains genomic contributions from e.g. *Malus sieversii, Malus sylvestris, Malus orientalis* and *Malus baccata*^*1*^. *M. baccata* is native to Asia and is used for breeding cultivars and rootstocks due to its distinct cold hardiness and disease resistance^2^. Breeding of new apple varieties is a complex process that takes several years^3^. Sequencing the genomes of apple genotypes provides insights into genetic diversity, evolutionary history, and genotype-phenotype relationships. The availability of whole genome sequences facilitates the development of marker-assisted selection (MAS), which aids breeding by enabling more efficient and targeted selection strategies. Recently, many genomes from various apple cultivars and individual accessions of different *Malus* species have been sequenced and published^4^, including that of *M. baccata*^*2*^. However, until now, no genome sequence has been available for *M. baccata* ‘Jackii’, an ornamental genotype collected in 1905 by J. G. Jack in Seoul^5^, which exhibits resistance to several fungal diseases such as apple scab (*Venturia inaequalis*)^6^, powdery mildew (*Podosphaera leucotricha*)^7^ and apple blotch (*Diplocarpon coronariae)*^8^. In addition, *M. baccata* ‘Jackii’ is resistant to fire blight^9,10^, caused by the Gram-negative bacterium *Erwinia amylovora*, which is one of the most destructive bacterial diseases affecting the genus *Malus*^*11*^. Breeding resistant apple cultivars is a promising and desirable strategy against fire blight and would require pyramiding the different resistance gene (R-gene) candidates due to the fact that resistance is strain-dependent, with some R-gene donors already overcome by virulent strains of *E. amylovora*^*11,12*^. The first fire blight R-gene to be isolated and functionally characterized using a transgenic approach was *FB_Mr5*, which underlies the resistance QTL region at the top of chromosome 3 of *Malus* × *robusta* 5^13,14^. *FB_Mr5* encodes a CC-NBS-LRR resistance protein that interacts with the cysteine protease AvrRpt2_*Ea*_ from *E. amylovora*, demonstrating a gene-for-gene relationship^9,13–16^. Moreover, sequence analysis of *avrRpt2*_*Ea*_ from various *E. amylovora* strains, along with inoculation experiments using *avrRpt2*_*Ea*_ knock-out mutants, revealed that *FB_Mr5*-mediated resistance in *Malus* × *robusta* 5 is lost when inoculated with knockout mutants and strains carrying a cysteine-to-serine substitution at amino acid position 156 in the bacterial effector *avrRpt2*_*Ea*_. Furthermore, it was shown that fire blight resistance in *M. baccata* ‘Jackii’ likely functions in a similar manner to that of *Malus* × *robusta* 5^9,16^. Additionally, a closely related homologue of *FB_Mr5* was identified in *M. baccata* ‘Jackii’^17^. However, as it has been demonstrated that even a single amino acid substitution in specific regions of FB_Mr5 can have a significant impact and trigger autoactivity^18^, it is important to know the exact sequences of resistance genes and whether there are other candidate genes at an R-gene locus that show a very high sequence similarity. In this study, we generated a high-quality, haplotype-resolved genome assembly and annotation of *M. baccata* ‘Jackii’ by combining PacBio HiFi sequencing with Hi-C and mRNA sequencing. Targeted genotyping-by-sequencing (tGBS)^19^ data of an F_1_ biparental population derived from an ‘Idared’ × *M. baccata* ‘Jackii’ cross were generated, and single-nucleotide polymorphisms (SNPs) were identified by mapping the sequences to the newly assembled genome. By additionally mapping the sequences of the tGBS analysis to the HFTH1 reference genome, chromosome names were assigned based on the HFTH1 assembly^20^. The genome assembly and annotation presented here for this multi-resistant genotype are of significant value and can be utilised directly for various resistance analyses and other applications.

## Methods

### Sampling, DNA and RNA extraction

Leaves from *M. baccata* ‘Jackii’ were collected at the Fruit Genebank of the Julius Kühn-Institut (JKI) in Dresden-Pillnitz, Germany. The QIAGEN Genomic-tip 20/G kit (QIAGEN, Hilden, Germany) was used for DNA extraction, while the RNAprep Pure Plant Plus Kit (Tiangen, Beijing, China) was employed for RNA extraction, with both procedures conducted in accordance with the manufacturer’s protocols.

### Genome size estimation using Illumina sequencing

Genomic DNA from diploid *M. baccata* ‘Jackii’ was sequenced using the Illumina NovaSeq 6000 platform (Illumina, Inc., San Diego, CA, USA) with a paired-end read length of 150 bp and a 350 bp sequencing library. Subsequently, the reads were quality-filtered (polyG tails trimmed, minimum length ≥ 100 bp, average read quality ≥ Q20, homopolymer filter ≤ 10% consecutive identical bases, ≤ 50% of bases with Q < 10). After filtering, a total of 80.14 GB of sequencing data was obtained, corresponding to an estimated sequencing depth of ~135.19×. The GC content was 37.22%, with Q20 exceeding 97.85% and Q30 surpassing 93.87%. Genome analysis was conducted using the Jellyfish 2.1.4^21^ and GenomeScope 2.0^22^ software. The *k*-mer distribution map with *k* = 19 was generated, the haploid genome size of *M. baccata* was estimated to be 592,786,100 bp and the heterozygosity rate was calculated to be ~1.57% (Fig. 1). The repeat sequence content was estimated at ~52.78%.

**Fig. 1.**
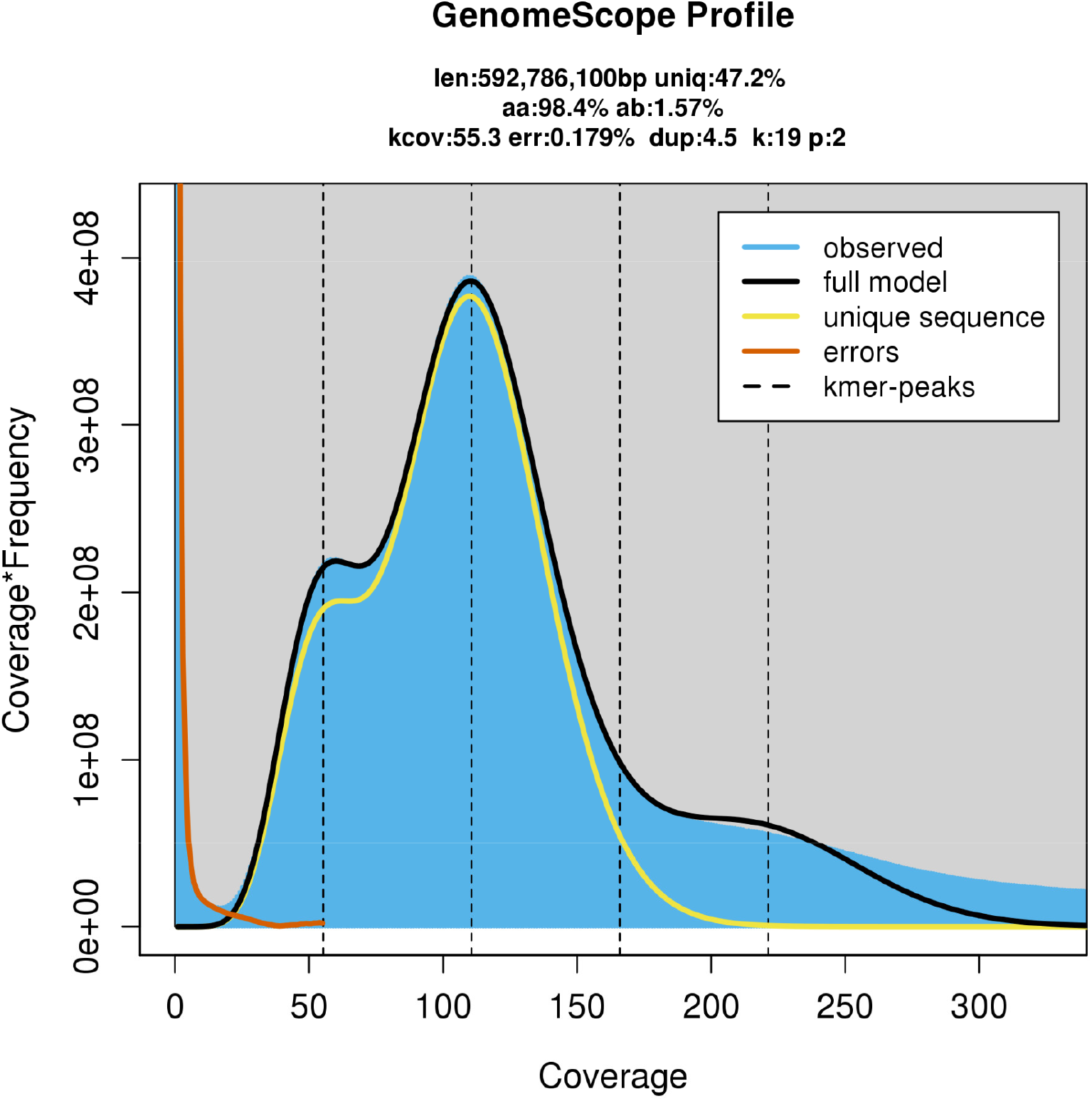
*K*-mer distribution map (*k*=19) of *Malus baccata* ‘Jackii’.

### Haplotype-resolved genome assembly with PacBio and Hi-C

DNA from *M. baccata* ‘Jackii’ was used for PacBio HiFi sequencing on the PacBio Revio platform (Pacific Biosciences, Menlo Park, CA, USA) following the standard protocol. The DNA molecules were sequenced in zero-mode waveguides (ZMWs) over multiple cycles, and repeated subreads were combined to generate highly accurate, self-corrected circular consensus sequencing (CCS) reads. In total, 6,149,970 CCS reads were produced, yielding 101,342,658,289 bp of sequence data. The average CCS read length was 16,479 bp, with an N50 of 16,808 bp, and the longest CCS read measured 61,744 bp.

In addition, an *in situ* Hi-C experiment was conducted^23^. To preserve DNA-DNA interactions and maintain the 3D genome structure, cross-linking was performed with formaldehyde and subsequently DNA was digested with the *Hin*dIII restriction enzyme generating sticky ends that were filled in with biotin-labeled nucleotides. Blunt-end ligation was performed to form circular structures. After reversing the cross-linking, the DNA was purified and sheared into fragments between 300-700 bp. Biotinylated junctions were isolated using streptavidin beads, and purified fragments were utilised for library preparation and sequencing on an Illumina NovaSeq 6000 PE150 (Illumina, Inc., San Diego, CA, USA). This process generated 689,199,106 read pairs, corresponding to 206,283,188,268 bp of sequence data, with an average GC content of 39.54%, a Q20 value of 97.98%, and a Q30 value of 94.70%. The Hi-C data were processed using HiC-Pro v2.10.0^24^, and the paired-end reads were aligned using BWA (v0.7.10-r789; mode: aln; default settings)^25^ to the preliminary assemblies of haplotype 1 and haplotype 2 derived from the CCS reads generated with Hifiasm (v0.19.9-r616)^26^. Of the total 1,378,398,212 reads, 1,174,709,293 and 1,175,354,792 reads were mapped to the preliminary assemblies of haplotype 1 and 2, respectively. Among these, 538,129,731 and 538,891,702 reads were uniquely mapped, resulting in 196,878,252 (36.59%) and 196,590,135 (36.48%) valid interaction pairs for haplotype 1 and 2, respectively. The preliminary assembly was segmented into 50-kb fragments and reassembled using the Hi-C data. Haplotype-resolved chromosome assembly was performed with LACHESIS^27^. A summary of the Hi-C-based haplotype-resolved genome assembly is presented in Table 1.

**Table 1.** Summary of the Hi-C-based haplotype-resolved genome assembly (only scaffolds > 1 kb were included).

| Haplotype | 1 | 2 |
| --- | --- | --- |
| Number of scaffolds | 719 | 272 |
| Total length of scaffolds | 654,629,689 | 637,559,422 |
| Scaffold N50 length | 35,776,412 | 36,609,931 |
| Scaffold N90 length | 30,157,701 | 30,099,248 |
| Maximum length of scaffold | 51,774,311 | 51,816,499 |
| Number of contigs | 740 | 292 |
| Total length of contigs | 654,627,589 | 637,557,422 |
| Contig N50 length | 30,388,229 | 30,684,037 |
| Contig N90 length | 9,863,630 | 11,516,422 |
| Maximum length of the contigs | 51,774,311 | 51,816,499 |
| GC content (%) | 38.14 | 38.02 |
| Gap numbers | 21 | 20 |

The assembled genome sequence was cut into 300 kb bins, and the signal intensity between corresponding bins was visualized as a heatmap in Fig. 2. The signal intensity was stronger within the 17 chromosome groups than between them, indicating a high-quality genome assembly.

**Fig. 2.**
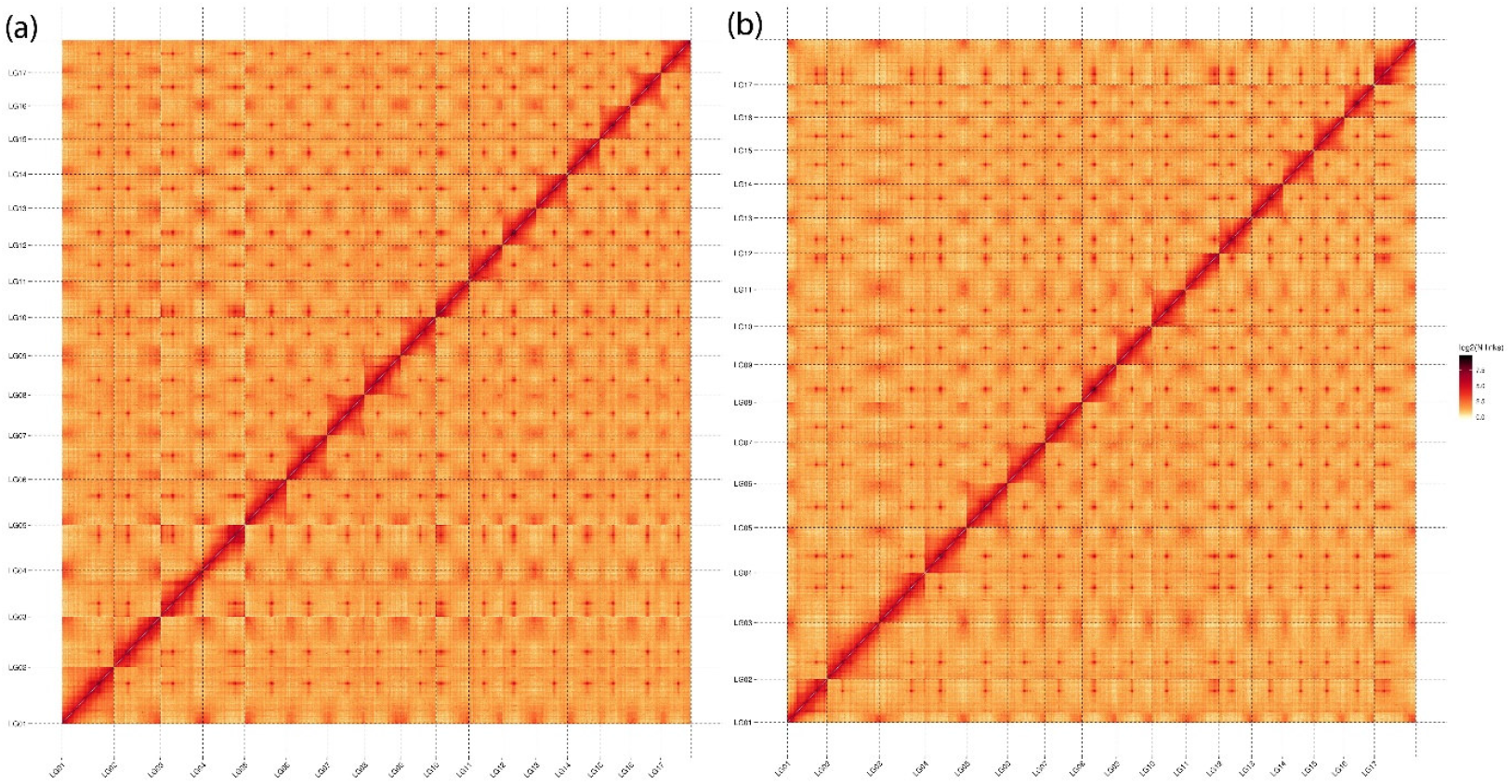
Heatmap of the assembled haplotype-resolved genomes. (a) Haplotype 1. (b) Haplotype.

### mRNA sequencing and genome annotation

The assembled genome was then used for genome annotation, with transposable element prediction performed using the following programs and databases: RepeatModeler2 v2.0.1^28^, RECON v1.0.8^29^, RepeatScout v1.0.6^30^, LTR_retriever v2.8^31^, LTRharvest v1.5.9^32^, LTR_FINDER v1.1^33^, RepeatMasker v4.1.0^34^, Repbase v19.06^35^, REXdb v3.0^36^ and Dfam v3.2^37^. Tandem repeats were predicted using the Microsatellite identification tool (MISA v2.1)^38^ and the Tandem Repeat Finder (TRF, v409)^39^. The results of the aforementioned analyses are presented in Tables 2 and 3.

**Table 2.** Summary of predicted transposable elements.

| **Type of transposable element** | **Number of predicted repetitive sequences (HT1/HT2)** | **Sequence length in bp (HT1/HT2)** | **Percentage of genome (HT1/HT2)** |
| --- | --- | --- | --- |
| Class I: DIRS | 2/3 | 122/169 | 0.00/0.00 |
| Class I: LINE | 33,232/29,751 | 19,082,791/12,588,448 | 2.92/1.97 |
| Class I: LTR Caulimovirus | 1,167/1,511 | 1,680,429/1,567,889 | 0.26/0.25 |
| Class I: LTR Copia | 74,041/75,763 | 61,731,539/60,267,561 | 9.43/9.45 |
| Class I: LTR ERV | 4,421/4,361 | 404,792/335,617 | 0.06/0.05 |
| Class I: LTR Gypsy | 84,129/81,591 | 102,424,853/99,955,957 | 15.65/15.68 |
| Class I: LTR Ngaro | 285/300 | 21,356/23,580 | 0.00/0.00 |
| Class I: LTR Pao | 135/93 | 12,134/6,123 | 0.00/0.00 |
| Class I: LTR unknown | 176,052/180,631 | 78,416,294/81,106,872 | 11.98/12.72 |
| Class I: SINE | 16,693/16,433 | 2,778,885/2,248,211 | 0.42/0.35 |
| Total for Class I | 390,157/390,437 | 266,553,195/258,100,427 | 40.72/40.48 |
| Class II: Academ | 1/1 | 138/138 | 0.00/0.00 |
| Class II: CACTA | 2,613/3,019 | 1,102,211/983,375 | 0.17/0.15 |
| Class II: Crypton | 22/34 | 1,050/1,759 | 0.00/0.00 |
| Class II: Dada | 243/233 | 12,084/11,497 | 0.00/0.00 |
| Class II: Ginger | 35/33 | 1,712/1,407 | 0.00/0.00 |
| Class II: Helitron | 147,768/141,516 | 37,977,788/42,416,743 | 5.80/6.65 |
| Class II: IS3EU | 179/187 | 10,934/10,119 | 0.00/0.00 |
| Class II: Kolobok | 276/290 | 20,418/23,008 | 0.00/0.00 |
| Class II: Maverick | 156/241 | 13,930/22,666 | 0.00/0.00 |
| Class II: Merlin | 149/140 | 7,271/7,513 | 0.00/0.00 |
| Class II: Mutator | 461/541 | 29,174/38,009 | 0.00/0.01 |
| Class II: P | 75/81 | 5,351/4,763 | 0.00/0.00 |
| Class II: PIF-Harbinger | 5,776/5,750 | 3,113,051/3,104,728 | 0.48/0.49 |
| Class II: PiggyBac | 66/66 | 3,304/2,620 | 0.00/0.00 |
| Class II: Sola | 0/1 | 0/95 | 0.00/0.00 |
| Class II: Tc1-Mariner | 212/225 | 12,471/13,478 | 0.00/0.00 |
| Class II: unknown | 126,862/121,450 | 30,761,945/28,104,547 | 4.70/4.41 |
| Class II: Zisupton | 95/100 | 4,301/4,596 | 0.00/0.00 |
| Class II: hAT | 4,953/4,985 | 2,128,608/2,120,043 | 0.33/0.33 |
| Total for Class II | 289,942/278,893 | 75,205,741/76,871,104 | 11.49/12.06 |
| Unknown | 39/28 | 2,455/1,492 | 0.00/0.00 |
| Total | 680,138/669,358 | 341,761,391/334,973,023 | 52.21/52.54 |

HT: haplotype.

**Table 3.**
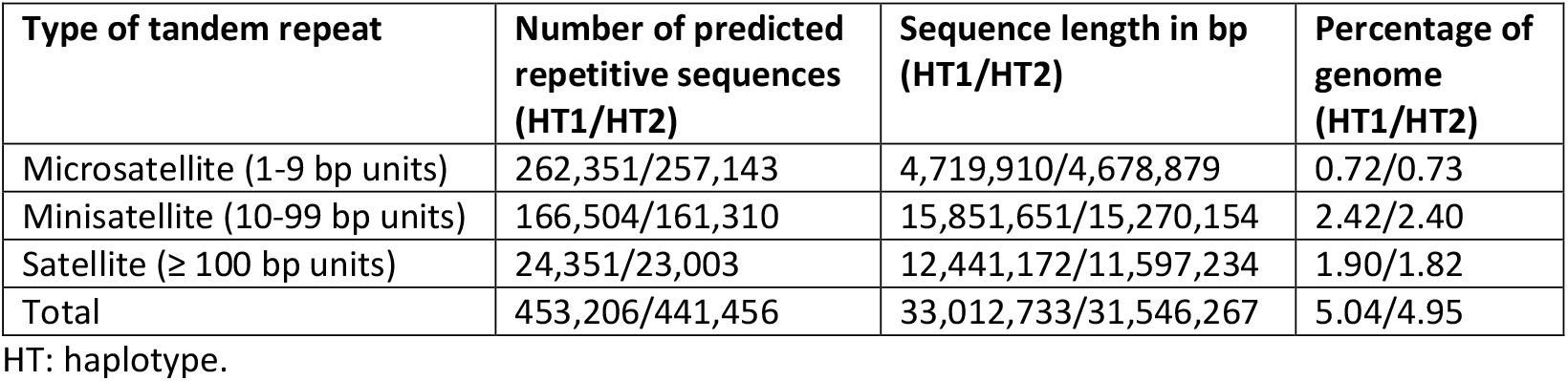
Summary of tandem repeat analysis.

The coding gene prediction was performed using three complementary approaches: *ab initio*, homology-based, and transcriptome-based methods (with and without reference genomes). *Ab initio* coding gene prediction was carried out using Augustus v2.4^40^ and SNAP (2006-07-28)^41^. Homology-based predictions were performed with GeMoMa v1.7^42^, and for transcriptome-based predictions, mRNA sequencing was performed on Illumina Novaseq 6000 platform (Illumina, Inc., San Diego, CA, USA) with a paired-end read length of 150 bp. This yielded a total of 40,279,131 reads (12.07 Gb), corresponding to 12,065,202,432 bp. The Q30 and Q20 values were 94.57% and 98.01%, respectively, and the GC content was 48.16%. Transcripts were predicted using HISAT v2.0.4^43^ and StringTie v1.2.3^44^ with different reference genomes^20,45–47^. Coding genes were then identified using GeneMarkS-T v5.1^48^. Additionally, transcripts were assembled without the use of a reference genome with Trinity v2.11^49^, and coding genes were predicted with PASA v2.0.2^50^. Finally, the genes predicted by the different methods were integrated using EVM v1.1.1^51^ and finalized by PASA v2.0.2^50^. The total number of coding genes predicted for haplotype 1 and 2 was 42,441 and 46,507, respectively. Detailed results are presented in Tables 4 and 5 and in Fig. 3.

**Table 4.**
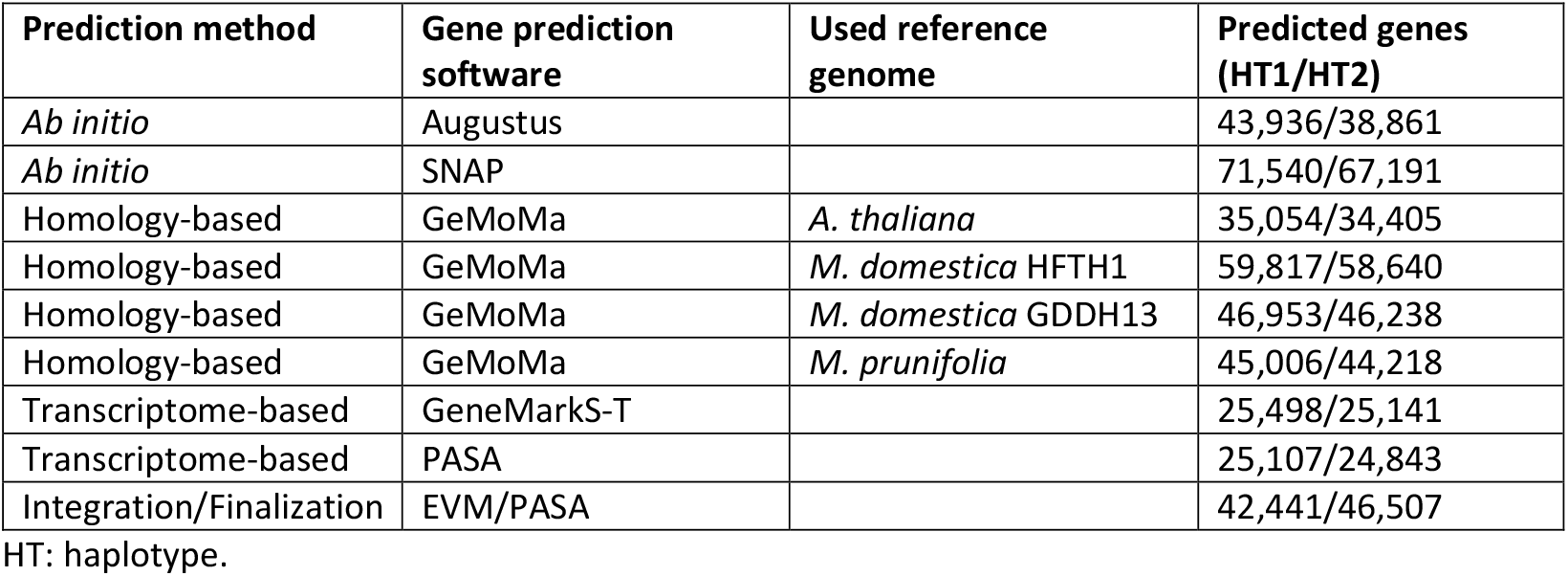
Predicted coding genes using different software tools.

| Prediction method | Gene prediction software | Used reference genome | Predicted genes (HT1/HT2) |
| --- | --- | --- | --- |
| *Ab initio* | Augustus |  | 43,936/38,861 |
| *Ab initio* | SNAP |  | 71,540/67,191 |
| Homology-based | GeMoMa | *A. thaliana* | 35,054/34,405 |
| Homology-based | GeMoMa | *M. domestica* HFTH1 | 59,817/58,640 |
| Homology-based | GeMoMa | *M. domestica* GDDH13 | 46,953/46,238 |
| Homology-based | GeMoMa | *M. prunifolia* | 45,006/44,218 |
| Transcriptome-based | GeneMarkS-T |  | 25,498/25,141 |
| Transcriptome-based | PASA |  | 25,107/24,843 |
| Integration/Finalization | EVM/PASA |  | 42,441/46,507 |

HT: haplotype.

**Table 5.** Statistics of predicted coding genes of *Malus baccata* ‘Jackii’.

| Haplotype | 1 | 2 |
| --- | --- | --- |
| Number of predicted genes | 42,441 | 46,507 |
| Total length of genes (bp) | 156,520,721 | 157,297,080 |
| Average gene length (bp) | 3,687.96 | 3,382.22 |
| Total length of exons (bp) | 69,643,903 | 71,153,751 |
| Number of exons | 234,373 | 238,393 |
| Average exons per gene | 5.52 | 5.13 |
| Total length of CDS (bp) | 59,033,112 | 60,656,154 |
| Total length of introns (bp) | 86,876,818 | 86,143,329 |
| Number of introns | 191,932 | 191,886 |
| Average introns per gene | 4.52 | 4.13 |

CDS: coding DNA sequence

**Fig. 3.**
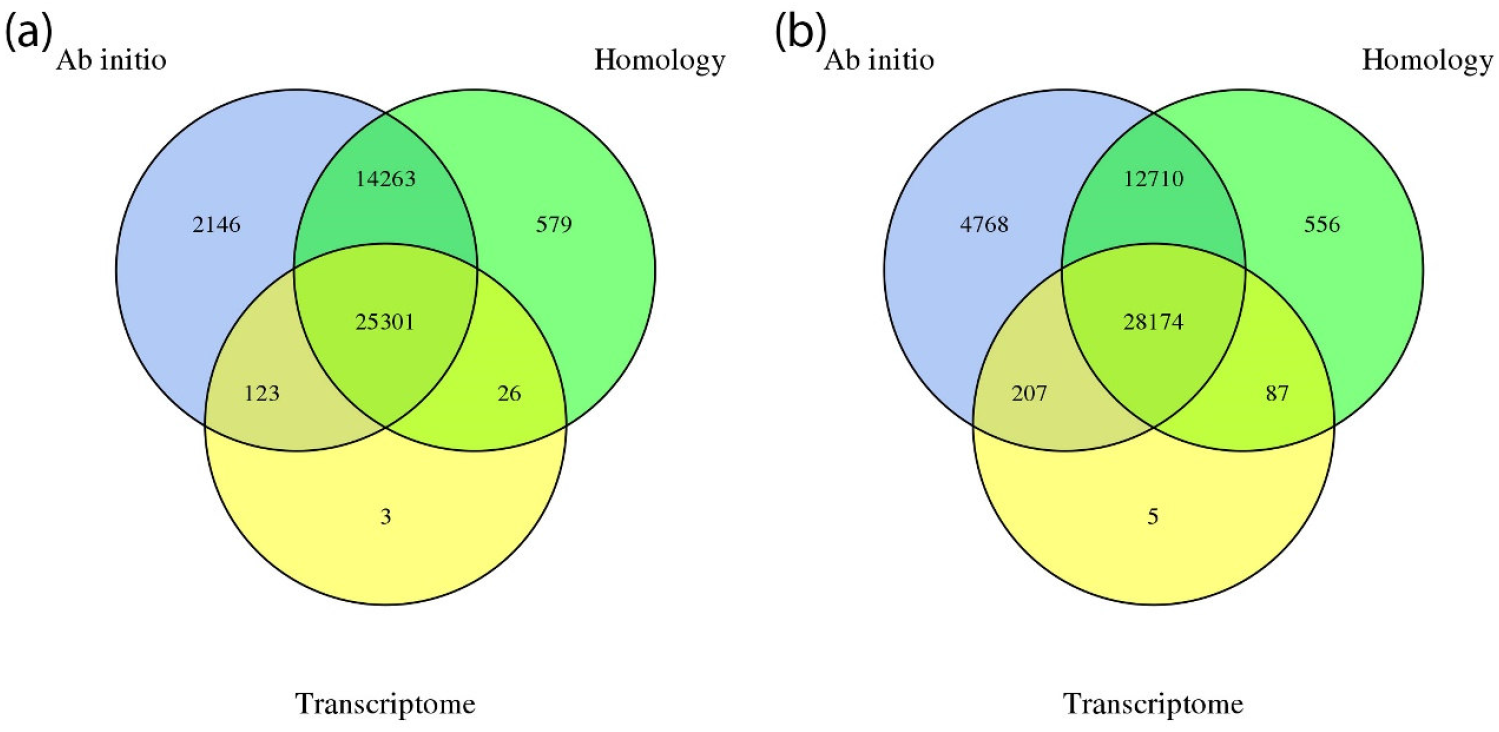
Venn diagram of integrated predicted coding genes. (a) Haplotype 1. (b) Haplotype 2. Created with Venny^52^.

Non-coding RNA prediction was conducted with tRNAscan-SE v1.3.1^53^ for tRNA, with barrnap v0.9^54^ based on Rfam v12.0^55^ for rRNA, with miRBase^56^ for miRNA, and with Infernal 1.1^57^ based on Rfam v12.0^55^ for snoRNA and snRNA. The results are summarised in Table 6.

**Table 6.**
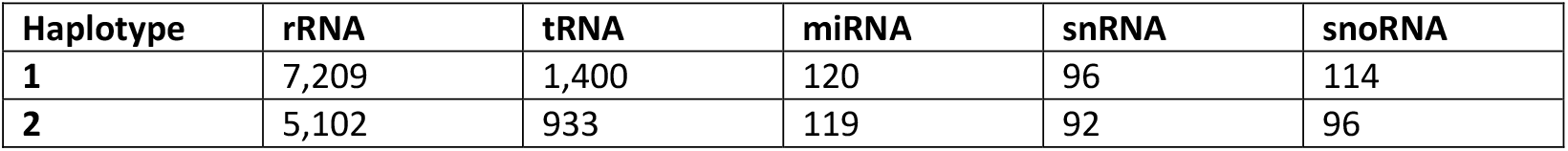
Predicted numbers of non-coding RNAs.

Homologous gene sequences without a complete gene locus were identified using GenBlastA v1.0.4^58^ and GeneWise v2.4.1^59^ was used to detect premature stop codons and frameshift mutations, resulting in the identification of 228 and 308 pseudogenes, respectively (Table 7).

**Table 7.** Predicted pseudogenes in *Malus baccata* ‘Jackii’.

| Haplotype | Number of predicted pseudogenes | Total length of predicted pseudogenes (bp) | Average length of pseudogenes (bp) |
| --- | --- | --- | --- |
| 1 | 228 | 975,909 | 4,280.30 |
| 2 | 308 | 1,282,971 | 4,165.49 |

The predicted coding genes were functionally annotated using multiple databases, including GenBank Non-Redundant (NR, 20200921), eggNOG 5.0^60^, Gene Ontology (GO, 20200615)^61,62^, Kyoto Encyclopedia of Genes and Genomes (KEGG, 20191220)^63^, SWISS-PROT and TrEMBL (202005)^64^, Pfam v33^65^ and eukaryotic orthologous groups (KOG, 20110125). Overall, more than 99.9% of coding genes were successfully annotated. The statistics on gene function annotation are presented in Table 8.

**Table 8.** Statistics of gene function annotation.

| Database | Number of annotated genes (HT1/HT2) | Percentage of annotated genes (HT1/HT2) |
| --- | --- | --- |
| GO | 33,072/33,956 | 77.92/73.01 |
| KEGG | 30,493/31,763 | 71.85/68.3 |
| KOG | 22,503/23,325 | 53.02/50.15 |
| Pfam | 34,025/35,128 | 80.17/75.53 |
| SWISS-PROT | 30,036/30,202 | 70.77/64.94 |
| TrEMBL | 42,402/46,449 | 99.91/99.88 |
| eggNOG | 34,189/35,057 | 80.56/75.38 |
| NR | 41,015/44,867 | 96.64/96.47 |
| Total | 42,408/46,461 | 99.92/99.9 |

HT: haplotype.

In the final step, InterProScan (5.34-73.0)^66^ was utilised for the prediction of motifs and domains. A total of 1,876 motifs and 45,003 domains were predicted in haplotype 1, and 1,820 motifs and 44,973 domains in haplotype 2.

### Chromosome assignment according to HFTH1

DNA of the F_1_ population (‘Idared’ × *M. baccata* ‘Jackii’), including the parental genotypes, was analysed using tGBS^19^ by Data2Bio (Ames, IA, USA). The restriction enzyme *Bsp*12861 was used and sequencing was carried out on an Illumina HiSeq X instrument (Illumina, Inc., San Diego, CA, USA). Polymorphic sites were first identified, and in a second step, final SNP calling was performed. In the initial step, individual sequence reads were scanned for low-quality regions (PHRED score ≤ 15), and quality-trimmed sequence reads were aligned to the genome sequences of haplotypes 1 and 2 of *M. baccata* ‘Jackii’ reported here using GSNAP^67^. Only confidently mapped reads with ≤ 2 mismatches per 36 bp and no more than 4 bases as tails per 75 bp that aligned to a single location were used for SNP identification, based on the following criteria: for homozygous SNPs, the most common allele had to be supported by at least 5 unique reads and 80% of all aligned reads. For heterozygous SNPs, each of the two most common alleles had to be supported by at least 5 unique reads and at least 30% of all aligned reads. For both homozygous and heterozygous SNPs, polymorphisms in the first and last 3 bp of each quality-trimmed read were ignored, and a PHRED base quality value of 20 (≤ 1% error rate) was set as the threshold for each polymorphic base. In the second step, SNPs were classified for tGBS genotyping as follows: a SNP was classified as homozygous if ≥ 5 reads supported the major allele and ≥ 90% of all reads at that site matched, and heterozygous if ≥ 2 reads supported each of two alleles, both alleles individually made up > 20%, and their combined reads were ≥ 5, covering ≥ 90% of all reads at that site. SNPs were then filtered based on a minimum calling rate of 50%, the allele number was set to 2, the number of genotypes ≥ 2, the minor allele frequency ≥ 10%, and the heterozygosity rate range between 0% and (2 × Frequency_allele1_ × Frequency_allele2_ + 20%). In a final step, imputation was used on chromosome-based SNPs that lacked a sufficient number of reads to make genotype calls using Beagle v5.4^68^ with 50 phasing iterations and default parameters. A total of 321,733 and 319,620 SNPs for haplotypes 1 and 2 were identified, with each site genotyped in at least 50% of the samples. SNP sequences from the tGBS data were then mapped to the HFTH1 genome sequence^20^ using BWA-MEM2^69^ on the JKI Galaxy Server (Galaxy v2.2.1+galaxy1)^70^ and the data presented in this study were assigned and oriented according to the HFTH1 reference^20^. A Circos plot^71^ illustrating key genomic features and alignments between the two haplotypes is shown in Fig. 4.

**Fig. 4.**
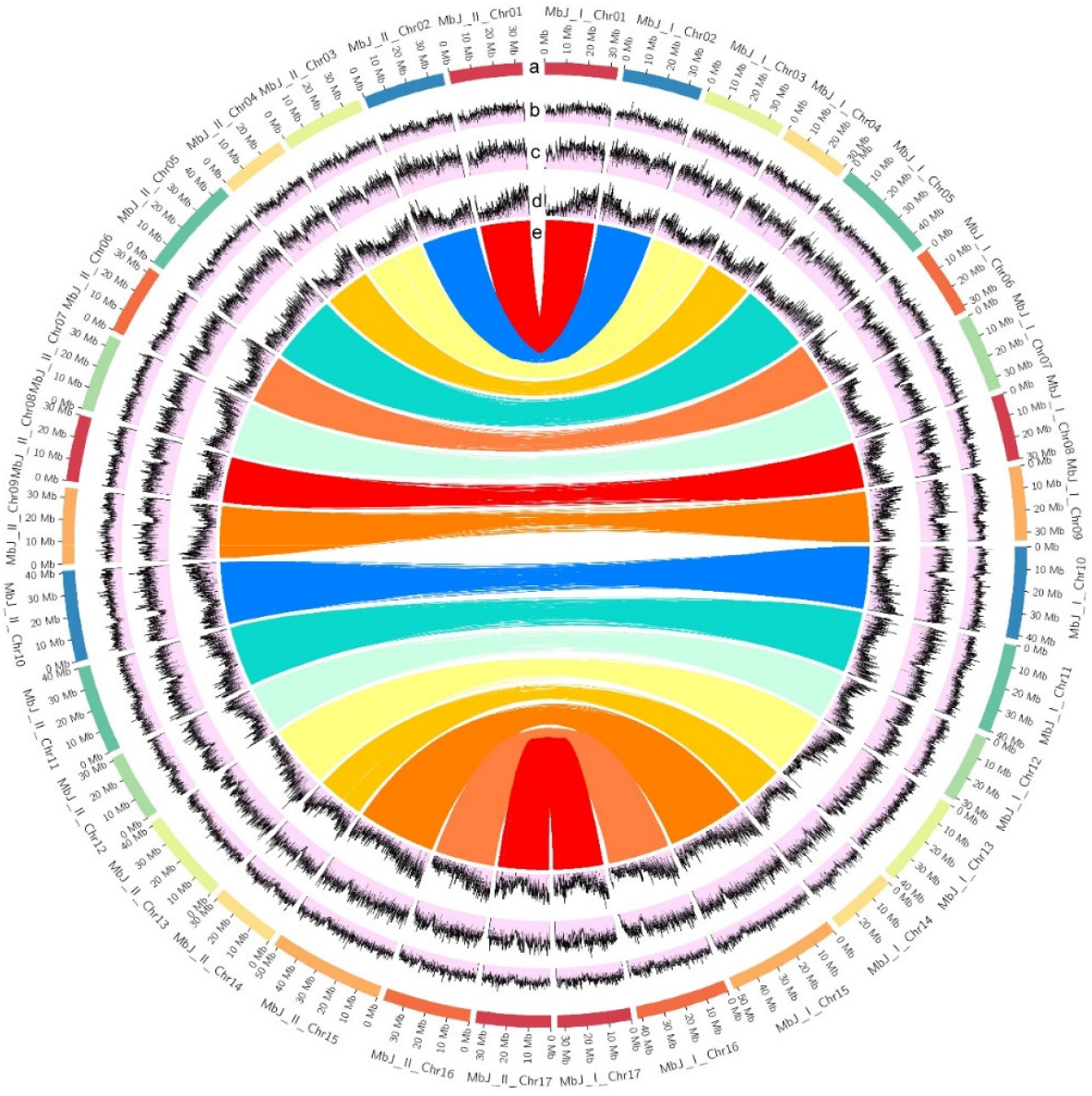
Circos plot of the *Malus baccata* ‘Jackii’ haplotype-resolved genome assembly. (a) Chromosome names and lengths in Mb. (b) Frequency of tandem repeats in 50 kb windows. (c) Frequency of transposable elements in 50 kb windows. (d) Frequency of genes in 50 kb windows. (e) Sequences from the tGBS analysis of haplotype 1 mapped onto haplotype 2. kb: kilobase, Mb: million base pairs, tGBS: targeted genotyping-by-sequencing.

### Data records

The raw data and assembled sequences and annotations can be accessed from the European Nucleotide Archive (ENA) under the BioProject accession number PRJEB89942 and study number ERP172974.

### Technical validation

To assess the completeness of the genome assembly, a BUSCO analysis with BUSCO v4.0^72^ was performed using the Embryophyta database containing 1,614 core genes. The results, presented in Table 9, demonstrate the high integrity of the haplotype-resolved genome assembly, with 97.58% and 97.52% of the core genes identified in haplotype 1 and 2, respectively.

**Table 9.** BUSCO analysis results using the Embryophyta database (1,614 core genes).

| Haplotype | Total number of identified core genes | Number of identified single-copy core genes | Number of identified duplicated core genes | Number of identified fragmented core genes | Number of missing core genes |
| --- | --- | --- | --- | --- | --- |
| **1** | 1,575 (97.58%) | 1,012 (62.70%) | 563 (34.88%) | 15 (0.93%) | 24 (1.49%) |
| **2** | 1,574 (97.52%) | 1,020 (63.20%) | 554 (34.42%) | 12 (0.74%) | 28 (1.73%) |

To functionally assess the quality of the genome assembly for downstream applications, the F_1_ population and both parents were grafted onto rootstock MM111 and up to five replicates per genotype were phenotyped for fire blight incidence i.e., length of shoot tip necrosis after artificial inoculation using *E. amylovora* strain *Ea*222, as described in Peil et al. (2007)^14^. Mean percent lesion length (PLL) was calculated for each F_1_ genotype by dividing the length of necrotic shoot by the total shoot length and averaging data from experiments conducted in 2024 and 2025. To identify potential associations between SNPs and fire blight incidence, each SNP with a minor allele frequency (MAF) ≥ 0.05 was analysed using the following procedure: genotypes were divided into two groups according to the observed allele. Using ‘Idared’ as a reference for the susceptible allele, phenotypic values of the two groups were used for a non-parametric Wilcoxon rank-sum test performed in R v4.4.2^73^. The genome-wide significance threshold was determined using Bonferroni correction as -log_10_(0.01/number of SNPs tested). A Manhattan plot was generated using the ggplot2 package^74^ and it showed a significant association of SNP markers at the top of chromosome 3 with the fire blight phenotypic data of the F_1_ progeny, shown for haplotype 2 in Fig. 5. Haplotype 1 produced a comparable plot (not shown). The sequence of the *FB_Mr5* homolog in *M. baccata* ‘Jackii’ (GenBank accession KT013244.1^17^) was found to be 100% identical over 4,164 bp to a region at the top of chromosome 3 of haplotype 2 between positions 598,337 and 602,501 bp, and this homolog is only 43,365 bp distant from the SNP marker with the highest -log_10_(p) value and the strongest association with fire blight resistance. This supports the accuracy of the haplotype-resolved genome assembly, the correctness of chromosome assignment and orientation, as *FB_Mr5* has been described to be located on the distal part of chromosome 3^14^, and the reliability and usability of the genomic data presented here for further analyses.

**Fig. 5.**
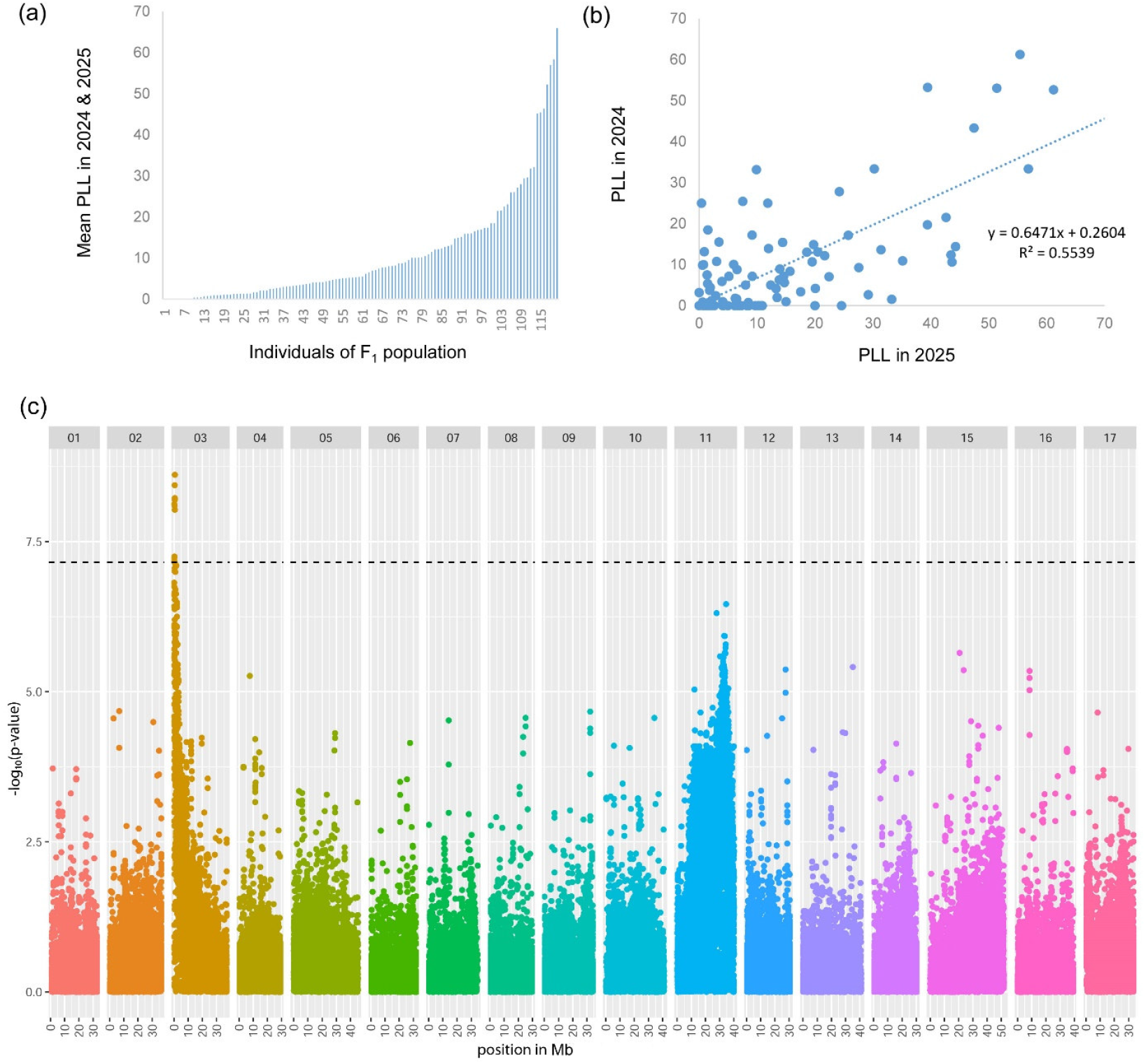
PLL in the F_1_ biparental (‘Idared’ × *M. baccata* ‘Jackii’) population after artificial fire blight inoculation. (a) Mean necrosis (%) across 2024 and 2025. (b) Correlation of necrosis between years across F_1_ genotypes. (c) Manhattan plot of the genome-wide association for mean PLL in the F_1_ population with newly generated SNP markers of haplotype 2. The dashed line indicates the Bonferroni-corrected significance threshold. Mb: million base pairs, PLL: percent lesion length, SNP: single-nucleotide polymorphism.

## Code availability

A custom R script was used to perform association mapping and is available from the corresponding author upon request.

## Acknowledgements

Parts of this work were supported by the Federal Ministry of Agriculture, Food and Regional Identity by decision of the German Bundestag (funding reference number: 281D108×21).

## Author information

### Authors and Affiliations

#### Julius Kühn-Institut (JKI) - Federal Research Centre for Cultivated Plants, Institute for Breeding Research on Fruit Crops, Dresden-Pillnitz, Germany

M. Pfeifer, O. F. Emeriewen, H. Flachowsky, M. Höfer, A. Peil & T. Wöhner

#### Institute of Plant Genetics, Department of Molecular Plant Breeding, Leibniz University Hannover, Hannover, Germany

M. Pfeifer

#### Julius Kühn-Institut (JKI) - Federal Research Centre for Cultivated Plants, Institute for Biosafety in Plant Biotechnology, Quedlinburg, Germany

J. Keilwagen & F.-S. Lim

#### Julius Kühn-Institut (JKI) - Federal Research Centre for Cultivated Plants, Institute for Resistance Research and Stress Tolerance, Quedlinburg, Germany

H. Zetzsche

## Contributions

Conception: Henryk Flachowsky, Andreas Peil and Thomas Wöhner. Strategy and design: Matthias Pfeifer, Ofere Francis Emeriewen and Thomas Wöhner. Analyses and writing: Matthias Pfeifer. Plant material: Monika Höfer and Andreas Peil. Fire blight phenotyping: Holger Zetzsche. Association mapping and analysis: Matthias Pfeifer, Ofere Francis Emeriewen, Jens Keilwagen, Fang-Shiang Lim and Thomas Wöhner. Data curation and upload: Jens Keilwagen and Fang-Shiang Lim. Funding: Thomas Wöhner. Supervision: Henryk Flachowsky and Thomas Wöhner. Revision: All authors. All authors read and approved the final manuscript.

### Corresponding authors

Correspondence to T. Wöhner.

## Ethics declarations

### Competing interests

The authors declare no competing interests.

## Notes

### Competing Interest Statement

The authors have declared no competing interest.

